# ExpressWeb: A Web application for clustering and visualization of expression data

**DOI:** 10.1101/625939

**Authors:** Bruno Savelli, Sylvain Picard, Christophe Roux, Christophe Dunand

## Abstract

The recent explosion of transcriptomics and proteomic data have resulted in vast amounts of datasets without connection and sometime too large to be easily analysed. Integration between datasets and analysis of an extracted datasets are limiting factors which need to be solved in order to make full use of the data and to connect data.

ExpressWeb is an online web tool that combines a Taylor clustering of expressed data sets to extract gene network with gene annotations to visualise the co-expression network. Data sets can become from personal or publically experiments. ExpressWeb allows to easily compute clustering on expression data and provides friendly and useful visualisation tools as heatmaps, graphs and networks, generating output images which can be used for scientific publications.

## Background

An increasing number of transcriptomic analyses are available for scientific community with a growing number of information. Since each experiment has been generated independently with a dedicated purpose, it is somewhere complex to compare experiment to one another. One of the perspectives resulting of the numerous available data is the analysis of expression profile per genes and the comparison between genes in order to produce expression gene cluster. Different web tools such as ATTED II (Aoki, et al., 2016), STRING database (Szklarczyk, et al., 2015), PlaNet (Ruprecht, et al., 2017), Planex (Yim, et al., 2013), CoP (Ogata et al., 2010) PlantExpress (Kudo, et al., 2017) ou COEXPRESSdb (Obayashi, et al., 2013) are available. The purpose of these programs is to represent expression gene cluster mainly for *Arabidopsis thaliana*.

By selecting the identifier of one or more genes, we can access to the network of genes co-expressed with the gene(s) of interest. These tools are useful for characterizing genes but do not allow the grouping of genes according to their expression profile and do not propose any statistical analysis. On the other hand, softwares used to cluster on large datasets without having to type code like HOMER (http://homer.ucsd.edu/homer/basicTutorial/clustering.html) or Cluster 3.0 (http://bonsai.hgc.jp/~mdehoon/software/cluster) are available. Unfortunately, these tools are not linked to a database, cannot incorporate annotation information, and do not allow tracing network coexpression.

Moreover, these resources display clustering based on already analysed data sets and it is not possible to include new expression data set in the co-expression analysis. In the particular case of model organisms such as *A. thaliana* the huge number of data available will create artefact clustering. The Taylor clustering proposed with selected data sets will allow bypassing this bias.

In order to perform efficient gene clustering on new data set together with gene annotation, we developed a flexible tool for analysis.

## Implementation

### Input data format

Data treated are expression values issuing from transcriptomic experiments (RNAseq, Fluidigm…). They are thus presented in the format of a file containing, for each gene, the expression values corresponding to the different replicates of the experimental conditions. They need to be in a CSV format including a header with the name of the experiment and for each gene its name and the expression values for each replicate/condition. A second file containing gene annotations can be added.

The number of genes tested ranged from a few hundred to a few tens of thousands depending on data sets and calculating capacity. Whereas local use of ExpressWeb is indeed possible for reduced data sets, an access to a computing cluster is necessary for large data sets (over 500-1000 genes). As the initial purpose of ExpressWeb was not to propose a statistical pipeline for transcriptomic analysis, the data will be processed as centering-reduction expression values. It is therefore for the user to provide a set of data that has been processed by differential analysis or any analysis of his choice.

### ExpressWeb: computational pipeline

Several languages have been used to develop ExpressWeb: HTML5 and CSS3 for the web interface, javascript for the dynamic tools (Highchart library to create dynamic Heatmap and the vis.js library to create network from the data), JSON (Kobayashi, et al., 2011) and JQuery for the use of DOM and Ajax in order to manage the data in dynamic manner. The language of the main script is PHP. Finally, once you have uploaded your values in the database, two scripts using R and Python languages will process it First, the R Script is used to compute hierarchical clustering on the up loaded values in order to build genes groups. Before computing the clustering, the values are scaled in order to have better results. The Varclus function is used, computing Hoeffding’s Distance between each gene and retrieving it into a similarity matrix. Then the Hclust function computes hierarchical clustering following the Ward’s algorithm creating a dendogramm. This dendogramm is cut to produce two types of groups, one with the threshold chosen by the user (Fig. 1A) and one with a lower threshold. The group definition and the scaled values are uploaded as MySQL database in order to optimize the re-use of the clustering results and to improve the performances during the analysis.

**Fig. 1.**
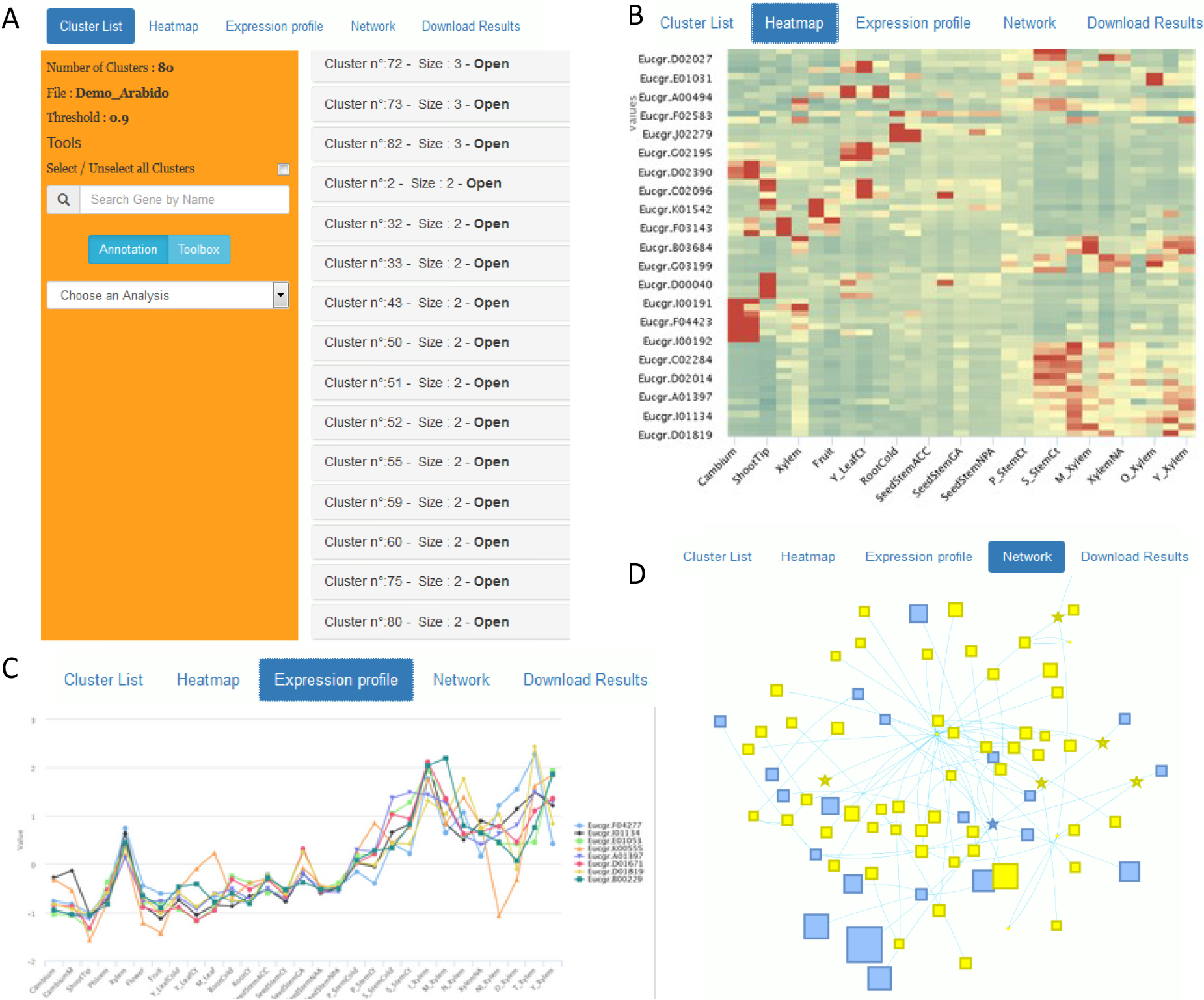
Screen shots of results following ExpressWeb analysis. **A**, List of the gene clusters. **B**, Heatmap of selected gene clusters. **C**, Expression profile of selected cluster, each gene profile can be removed from the vizualisation. **D**, Network obtained from all gene clusters, size of the icons is proportional to the numbers of genes per cluster.

The Python script will read the similarity matrix and the groups definition in order to create nodes and edges that will be used to build the network visualisation (Fig. 1D). Nodes are created for each gene. Each node contains information about the gene such as its name and its cluster. Then the nodes are connected by creating edges. Inside a group (low level clusters) the clusters are connected by computing average distance between each clusters and creating an edge between a cluster and the one having the best average distance. Inside a cluster, each gene is connected to its closest according to the similarity matrix except if those two are already connected and then the gene is connected to the second closest etc. Nodes and Edges are written in two JSON files that will be read using the vis.js library

### Web-server and standalone package

The idea of developing ExpressWeb came from our specific needs in analyzing selected sets of expression data. First ExpressWeb has been developed to be used in local with our own data set or with data set publically available, but rapidly a standalone package has been developed and userfriendly outputs such as heatmap, expression profile and network (Fig.1).

A demo version provides a user-friendly interface to ExpressWeb (http://polebio.lrsv.ups-tlse.fr/ExpressWeb).

## Conclusions

ExpressWeb provides a powerful web tool to perform a Taylor clustering analysis of personal or publically available experiments. It extracts gene network with gene annotations to visualise the co-expression network. Data sets can become from personal or publically experiments. ExpressWeb allows to easily compute clustering on expression data and provides friendly and useful visualisation tools as heatmaps, graphs and networks, generating output images which can be used for scientific publications.

## Availability and requirements

home page: Web-implemented version and source code are freely available at https://github.com/PeroxiBase/ExpressWeb and demo version can be tested at http://polebio.lrsv.ups-tlse.fr/ExpressWeb/.

## Acknowledgements

The authors thank Catherine Mathé for the scientific discussion and the Toulouse Midi-Pyrénées bioinformatics platform (Bioinfo Genotoul) for hosted the Demo version of ExpressWeb.

## Availability of data and materials

A detailed ExpressWeb documentation are available from github page

## Authors’ contributions

BS/CD conceived of the initial idea of ExpressWeb, BS/SP designed and programmed ExpressWeb, CR evaluated and provided ideas for ExpressWeb, and BS/CR/CD wrote the manuscript. All authors read and approved the final version.

